# HapIso: An Accurate Method for the Haplotype-Specific Isoforms Reconstruction from Long Single-Molecule Reads

**DOI:** 10.1101/050906

**Authors:** Serghei Mangul, Harry (Taegyun) Yang, Farhad Hormozdiari, Elizabeth Tseng, Alex Zelikovsky, Eleazar Eskin

## Abstract

Sequencing of RNA provides the possibility to study an individual’s transcriptome landscape and determine allelic expression ratios. Single-molecule protocols generate multi-kilobase reads longer than most transcripts allowing sequencing of complete haplotype isoforms. This allows partitioning the reads into two parental haplotypes. While the read length of the single-molecule protocols is long, the relatively high error rate limits the ability to accurately detect the genetic variants and assemble them into the haplotype-specific isoforms. In this paper, we present HapIso (**H**aplotype-specific **I**soform Reconstruction), a method able to tolerate the relatively high error-rate of the single-molecule platform and partition the isoform reads into the parental alleles. Phasing the reads according to the allele of origin allows our method to efficiently distinguish between the read errors and the true biological mutations. HapIso uses a k-means clustering algorithm aiming to group the reads into two meaningful clusters maximizing the similarity of the reads within cluster and minimizing the similarity of the reads from different clusters. Each cluster corresponds to a parental haplotype. We use family pedigree information to evaluate our approach. Experimental validation suggests that HapIso is able to tolerate the relatively high error-rate and accurately partition the reads into the parental alleles of the isoform transcripts. Furthermore, our method is the first method able to reconstruct the haplotype-specific isoforms from long single-molecule reads.

The open source Python implementation of HapIso is freely available for download at https://github.com/smangul1/HapIso/

## 1 Introduction

Advances in the RNA sequencing technologies and the ability to generate deep coverage data in the form of millions of reads provide an exceptional opportunity to study the functional implications of the genetic variability [4], [16], [17]. RNA-Seq has become a technology of choice for gene expression studies, rapidly replacing microarray approaches [20]. RNA-Seq provides sequence information, which aids in the discovery of genetic variants and alternatively spliced isoforms within the transcripts. RNA-Seq has the potential to quantify the relative expression of two alleles in the same individual and determine the genes subject to differential expression between the two alleles. Comparison of the relative expression of two alleles in the same individual as a phenotype influenced by the cis-acting genetic variants helps determine the cis-acting nature of the individual polymorphism [23].

There are three major difficulties in current approaches to identify allele specific expression using RNA-Seq data. First, short read protocols [15] cut genetic material into small fragments and destroy the linkage between genetic variants. Short reads obtained from the fragments are well suited to access the allele-specific expression on the single variant level. However, complexity of the higher eukaryotic genomes makes it hard to phase the individual variants into the full-length parental haplotypes (haplotype-specific isoforms). A common technique to assess the allele-specific expression (ASE) is to count the number of reads with the reference allele and the number of reads with alternate allele. However, this approach works on individual variant level and is not well suited to determine the allele-specific expression on the isoform level. Second, mapping the short reads onto the reference genome introduces a significant bias toward higher mapping rates of the reference allele at the heterozygous loci. Masking known loci in the genome does not completely remove the inherent bias [6]. Aside from the allele-specific expression, mapping biases may affect the QTL mapping and the discovery of new sequence variants. Third, the high sequence similarity between alternatively spliced variants of the same gene results in a significant number of short reads to align in multiple places of the reference transcriptome [9].

Multi-kilobase reads generated by single-molecule protocols [8] are within the size distribution of most transcripts and allow the sequencing of full-length haplotype isoforms in a single pass [14]. The reads cover multiple genomic variants across the gene, eliminating the necessity to phase the individual variants into the isoform haplotypes. Additionally, the extended length of the reads makes it simple to map the reads uniquely and eliminate the read-mapping biases. However, the relatively high error rates of the single-molecule protocols limit the application of the long single-molecule protocol to studies of the allele specific variants. There are currently no methods able to accurately detect the genetic variants from the long single-molecule RNA-Seq data and connect them to haplotype-specific isoforms.

In this paper, we present HapIso (**H**aplotype-specific **I**soform Reconstruction), a comprehensive method for the accurate reconstruction of the haplotype-specific isoforms of a diploid cell that uses the splice mapping of the long single-molecule reads and partitions the reads into parental haplotypes. The single molecule reads entirely span the RNA transcripts and bridge the single nucleotide variation (SNV) loci across a single gene. Our method starts with mapping the reads onto the reference genome. Aligned reads are partitioned into the genes as a clusters of overlapping reads. To overcome gapped coverage and splicing structures of the gene, the haplotype reconstruction procedure is applied independently for regions of contiguous coverage defined as *transcribed segments*. Restricted reads from the transcribed regions are partitioned into two local clusters using the 2-mean clustering. Using the linkage provided by the long single-molecule reads, we connect the local clusters into two global clusters. An error-correction protocol is applied for the reads from the same cluster. To our knowledge, our method is the first method able to reconstruct the haplotype-specific isoforms from long single-molecule reads. We applied HapIso to publicly available single-molecule RNA-Seq data from the GM12878 cell line [18]. Circular-consensus (CCS) single-molecule reads were generated by Pacific Biosciences platform [8]. Parental information (GM12891 and GM12892 cell lines) is used to validate the accuracy of the isoform haplotype reconstruction (i.e. assignment of RNA molecules to the allele of origin). We use short read RNA-Seq data for the GM12878 sample to validate the detected SNVs using a different sequencing platform (short reads). Our method discovered novel SNVs in the regions that were previously unreachable by the short read protocols.

Discriminating the long reads into the parental haplotypes allows to accurately calculate allele-specific gene expression and determine imprinted genes [7], [19]. Additionally it has a potential to improve detection of the effect of cis-and trans-regulatory changes on the gene expression regulation [5], [21]. Long reads allow to access the genetic variation in the regions previously unreachable by the short read protocols providing new insights into the disease sustainability.

## 2 Methods

### 2.1 Overview

Very similarly to the genome-wide haplotype assembly problem, the problem of haplotype-specific isoform assembly aims to infer two parental haplotypes given the collection of the reads [1], [12]. While those problems are related, the allele expression ratio between RNA haplotypes is a priori unknown and may be significantly different from 1:1. An additional difference is due to the RNA-Seq gapped alignment profile and alternative splice structures of the gene. Overall, the problem of reconstruction of the haplotype-specific isoforms of a diploid transcriptome represents a separate problem requiring novel computational approaches.

We apply a single-molecule read protocol to study the allele-specific differences in the haploid transcriptome (Figure 1). The single molecule protocol skips the amplification step and directly sequences the poly (A) selected RNA molecules. The reads generated by the protocol entirely span the RNA transcripts bridging the single nucleotide variation (SNV) loci across a single gene.

**Fig. 1.**
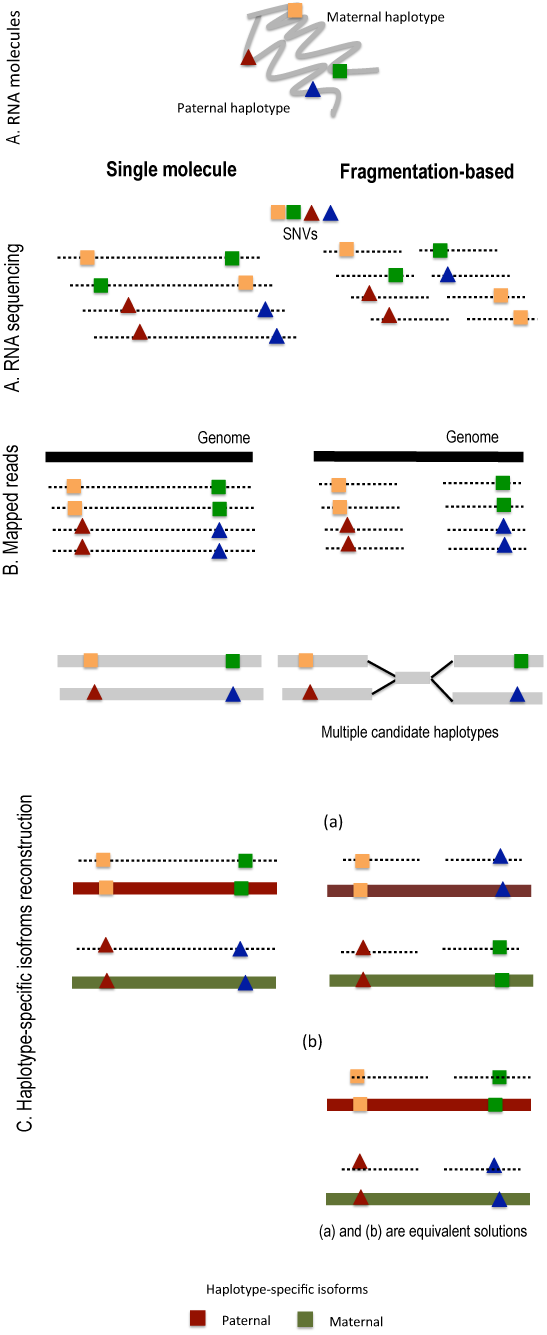
Overview of long single-molecule protocol. (A) Unamplifled, polyA-selected RNA molecules are sequenced by the single-molecule protocol able to entirely span the RNA transcripts to produce long single-molecule reads. The fragmentation-based protocols shred the amplified and poly(A) selected RNA into short fragments appropriately sized for sequencing. Short reads destroys the linkage between the SNVs. (B) Reads are mapped onto the reference genome. (C) SNVs are assembled into the two parental haplotype isoforms.

We introduce a method able to reconstruct the haploid transcriptome of a diploid organism from long single-molecule reads (Figure 2). This method is able to tolerate the relatively high error-rate of the single-molecule sequencing and to partition the reads into the parental alleles of the isoform transcript. The errors in the long single-molecule reads typically are predominantly one-base deletions and insertions [3]. Both insertions and deletions are corrected through the alignment with the reference genome. The remaining mismatch errors are further passed to the downstream analysis.

**Fig. 2.**
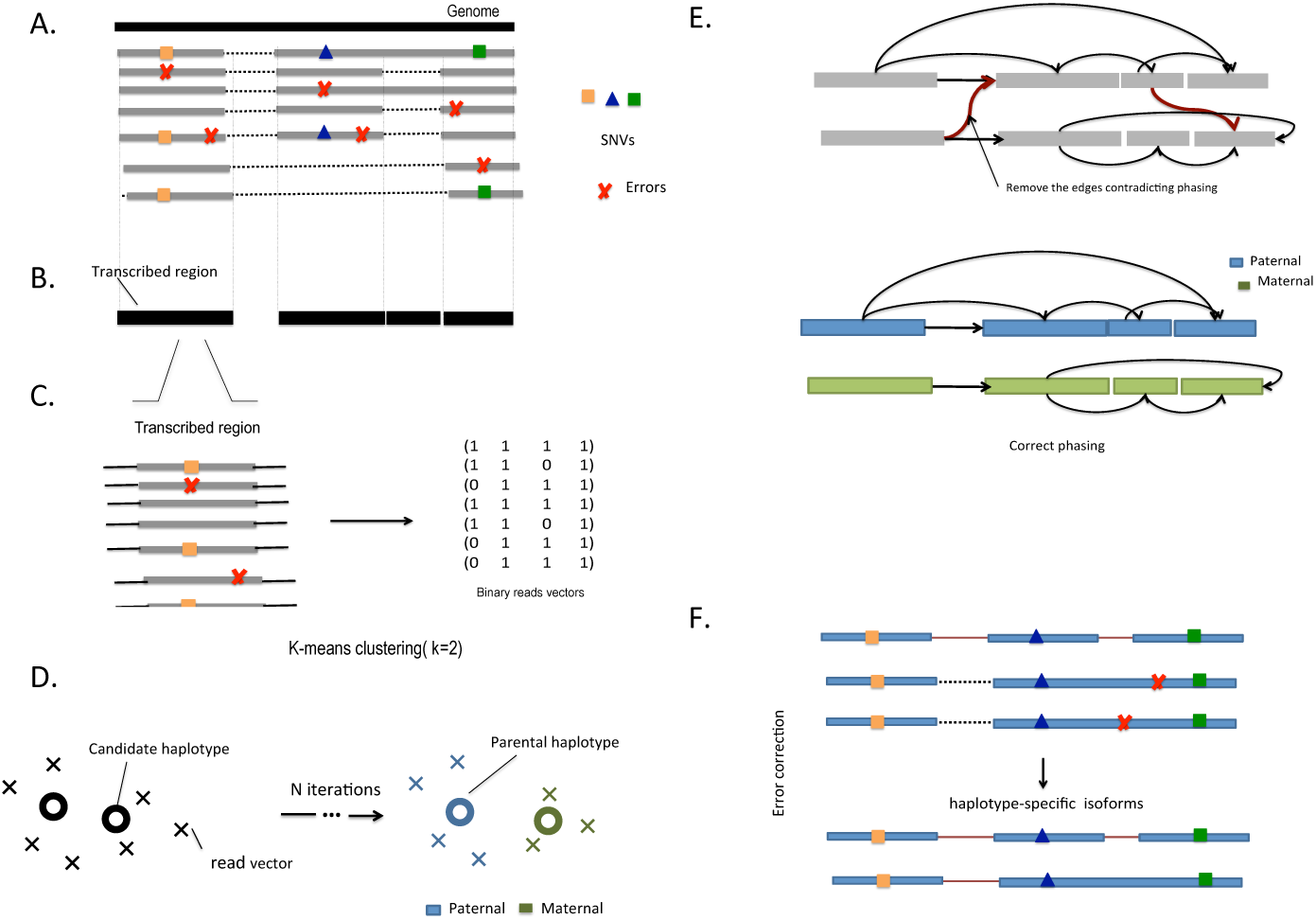
Overview of HapIso. (A) The algorithm takes long single-molecule reads that have been mapped to the reference genome as an input. (B) The transcribed segments are identified as contiguous regions of equivalently covered positions. (C) Aligned nucleotides of the transcribed segment are condensed into the binary matrix whose width equals the number of variable positions. The entry “1” corresponds to the position with the observed mismatch, the entry is encoded as “0” if it matches the reference allele. (D) Reads restricted to the transcribed segment (rows of the binary matrix) are partitioned into two clusters, using the 2-means clustering algorithm. Each cluster corresponds to a local haplotype. (E) The segment graph is constructed to incorporate the linkage between the alleles. The edges of the graph connect the local haplotypes. The minimum number of corrections to the graph is applied to partition the graph into two independent components corresponding to full-length parental gene haplotypes. (F) An error-correction protocol is applied for the reads from the same cluster. The protocol corrects the sequencing errors and produce corrected haplotype-specific isoforms.

Our method starts with mapping the reads onto the reference genome (Figure 2.A). Long reads allow us to identify the unique placement of the read (99.9% of the reads from GM12878 sample are mapped to a single location in the genome). The reads are partitioned into the genes as clusters of overlapping reads. The haplotype reconstruction procedure is applied independently for every gene. First, we identify the *transcribed segments* corresponding to contiguous regions of equivalently covered positions. Two positions are equivalently covered if any read covering one position also covers the other one.

To account for gapped coverage and splicing structures of the gene, we cluster the reads into two parental haplotypes for every transcribed segments independently (Figure 2.C). The clustering procedure first condenses the aligned nucleotides of the transcribed segment into a binary matrix with a width equal to the number of variable positions. The entry “1” corresponds to the position which mismatches the reference allele, while the entry is encoded as “0” if it matches the reference allele. We partition the rows (reads restricted to the transcribed segment) into two clusters, using the 2-means clustering algorithm (Figure 2.D). The result from the 2-means clustering partitions the restricted reads into local parental haplotypes. Using the linkage provided by the long single-molecule reads, we reconstruct the full-length gene haplotypes. We build the segment graph encoding the linkage between the alleles in form of edges (Figure 2.E). The minimum number of corrections to the graph is applied to partition the graph into two independent components corresponding to two parental haplotypes. The transcript reads are then grouped according to the allele of origin (Figure 2.F). An error-correction protocol is applied for the reads from the same cluster. The protocol corrects the sequencing errors and produces corrected haplotype-specific isoforms.

### 2.2 Single-Molecule RNA-Seq

We use publicly available single-molecule RNA-Seq data generated from the peripheral blood lymphocyte receptors for B-lymphoblastoid cell lines (GM12878 cell line) [18]. Additionally, we use parental long read RNA-Seq data from GM12891 and GM12892 cell lines to validate the accuracy of the proposed approach. Libraries were sequenced using the Pacific Bioscience platform [8] able to produce long single-molecule reads for all three samples in the trio. Unamplified, polyA-selected RNA was sequenced by the circular molecules. Circular-consensus (CCS) single-molecule read represent a multipass consensus sequence, where each base pair is covered on both strands at least once and the multiple low-quality base calls can be used to derive a high-quality calls.

### 2.3 Read Mapping

The first step of the analysis is to place sequencing reads onto the reference genome. Long read length provided by the single molecule protocol provides enough confidence to find unique position in the genome where the reads were generated from without using the existing gene structure annotation. The 715,902 CCS reads were aligned to the human reference genome (hg19) using the GMAP aligner, which maps cDNA sequences to a genome [22]. GMAP was originally designed to map both messenger RNAs (mRNAs) and expressed sequence tags (ESTs) onto to genome. The tool is able to tolerate high number of sequence errors in long cDNA sequences which makes it perfect fit for Pac Bio single-molecule platform.

GMAP is able to identify up to two placements in the genome for 99.6% reads. Only a small portion of those are mapped to two locations of the genome (1.6%). However in many case two placement in the genome have a evident differences thus making it easy to select the most preferable placement. In this way, vast majority of the CCS reads have single high-confidence mapping covering the entire exon-intron structure of an isoform transcript.

### 2.4 Haplotype-Specific Isoform Reconstruction

Having the reads spanning the full-length isoform transcripts, the problem of the haplotype-specific isoform reconstruction aims to discriminate the transcript reads into two parental haplotypes. The problem is equivalent to the read error-correction problem. If all the errors are corrected, the long reads provide the answer to the reconstruction problem, i.e. each non redundant read is the haplotype-specific isoform. Since the long reads are error prone, it is required to fix the errors or equivalently call single nucleotide variants (SNVs). Rather than phasing each isoform separately, it is preferable to agglomerate all the isoforms from a single gene and cluster the reads by haplotype of origin. All the reads from a single haplotype contain the same alleles of shared SNVs, thus all the differences between reads in shared transcribed segments should be corrected. We propose the following optimization formulation minimizing the number of errors.

#### Phasing Problem

Given a set of the long reads *R* corresponding to the transcripts from the same gene *g*. Partition the reads into two haplotype clusters such that the number of sequencing errors in the reads is minimized.

Typically, the errors in the long single-molecule reads are dominated by one-base insertions and deletions. Both are corrected through the alignment to the reference genome - insertions are deleted and deletions are imputed from the reference. The remaining mismatch errors are further passed to the downstream analysis.

Since long reads are uniquely aligned, the aligned reads are uniquely partitioned into clusters corresponding to the genes. Further, the haplotype are independently reconstructed for every gene. First we split the genes into *transcribed segments* corresponding to contiguous regions of equivalently covered positions. Two positions are equivalently covered if any read covering one position also covers the other one. To overcome gapped coverage and splicing structure of the gene, we cluster the reads into two parental haplotype for every transcribed segments independently. The clustering procedure first condenses the aligned nucleotides of the transcribed segment into the binary matrix whose width equals the number of polymorphic positions. The entry “1” corresponds to each position whose allele mismatches the reference and the entry is encoded as “0” if it matches the reference allele. We partition the rows (reads restricted to the transcribed segment) into two clusters, using the 2-means clustering algorithm. The clustering algorithm returns a set of centroids, one for each of the two clusters. An observation vector is classified with the cluster number or centroid index of the centroid closest to it. A vector *r* belongs to cluster *i* if it is closer to centroid *i* than any other centroids. If *r* belongs to *i*, centroid *i* is refereed as dominating centroid of *r*.

Given a set of observations (*r*_1_, *r*_2_, *r*_n_), where each observation is a binary read vector, k-means clustering aims to partition the n reads into two sets *S* = *S*_1_, *S*_2_ so as to minimize the distortion defined as

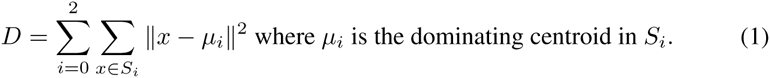

Each step of the k-means algorithm refines the choices of centroids to minimize distortion. The clustering algorithm uses change in distortion as stopping condition.

Haplotypes for each transcribed segment obtained by local read clustering are further linked using the long single-molecule reads as follows.

First, we build a graph *G* in which each vertex corresponds to a transcribed segment and two vertices are adjacent if they belong to the same read. Two transcribed segments *A* and *B* with pairs of haplotypes (*A*1, *A*2) and (*B*1, *B*2), respectively, can be either linked in parallel *A*1*B*1 and *A*2*B*2 or in cross *A*1*B*2 and *A*2*B*1. If among reads containing *A* and *B* there are more reads consistent with the parallel linkage than the reads consisting with the cross linkage, then the parallel linkage is more likely and vice versa. The larger skew between number of parallel and cross reads gives the higher confidence in the corresponding linkage. Therefore the weight of an edge between *A* and *B* in the graph *G* is set to

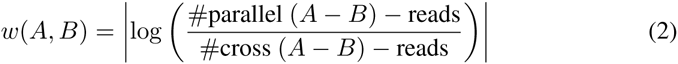

Then we find the maximum-weight spanning tree *T* of the graph *G* consisting of links with the highest confidence in the chosen (cross or parallel) linkage [2]. The tree *T* is split into two trees T1 and T2 uniquely identifying two global haplotypes for the corresponding gene as follows. Each vertex A of T is split into two vertices *A*1 and *A*2 corresponding to two haplotypes of the transcribed segment A. Each edge (*A*, *B*) of *T* with the positive weight *w*(*A*, *B*) is split into two edges (*A*1, *B*1) and (*A*2, *B*2) and each edge (*A*, *B*) with the negative weight is split into (*A*1, *B*2) and (*A*2, *B*1). Starting with *A*1 we traverse the tree *T*1 concatenating all haplotypes corresponding to its vertices into a single global haplotype. Similarly, starting with *A*2, we traverse the complementary tree *T*2 concatenating its haplotypes into the complimentary global haplotype.

Long reads are grouped according to the haplotype of origin. An error-correction protocol is applied for the reads from the same cluster. The protocol corrects the sequencing errors and produce corrected haplotype-specific isoforms.

Finally, the resulting two haplotypes are different in heterozygous loci allowing our method to determine the SNVs. Long reads provide one to one mapping between the reconstructed haplotypes and the isoform haplotypes. Reads counts are used to determine allelic expression of each haplotype copy of the isoform.

The absence of the systematic errors allows us to successfully correct the randomly distributed errors and accurately reconstruct the isoform haplotypes. Comparing to other approaches requiring trio family data, we are able to correct the error and reconstruct the parental haplotypes from sequencing data.

## 3 Results

### 3.1 HapIso Is Able To Accurately Reconstruct Haplotype-Specific Isoforms

We used trio family long single-molecule RNA-seq data to validate the reconstructed haplotype-specific isoforms. Single molecule RNA-Seq data was generated from the peripheral blood lymphocyte receptors for B-lymphoblastoid cell lines from a complete family trio composed of a father, a mother and a daughter. We reconstructed the haplotype-specific isoforms of each individual independently using HapIso method. Family pedigree information makes it possible for us to detect Mendelian inconsistencies in the data. We use reconstructed haplotypes to infer the heterozygous SNVs determined as position with non-identical alleles with at least 10x coverage.

According to Mendelian inheritance, one allele in the child should be inherited from one parent and another allele from the other parent. (Figure 3.A). Mendelian inconsistencies correspond to loci from the child with at least one allele not confirmed by parents (Figure 3.B). We also separately account for the missing alleles from the parental haplotypes due to insufficient expression or coverage of the alternative allele (Figure 3.C). Such loci are ambiguous since Mendelian consistency or inconsistency cannot be verified.

**Fig. 3.**
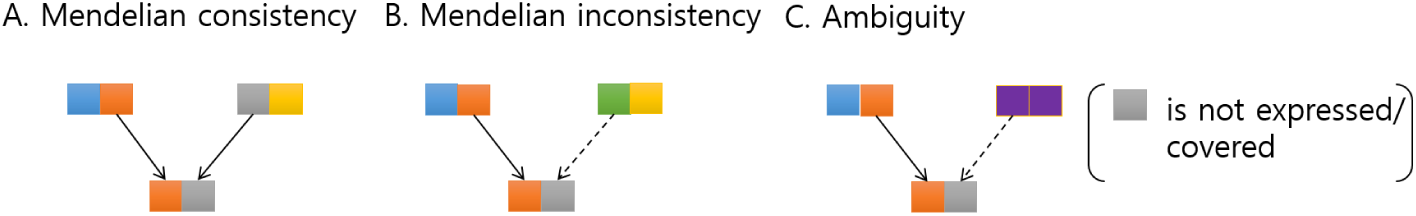
Allele inheritance model. (A) Mendelian consistency; Each parent provides one copy of the allele to a child; child inherits red allele from mother and a green allele from father. (B) Mendelian inconsistency; Child does not inherit green allele from any of the parents. (C) Ambiguity: Green allele is not expressed or covered in the right parent due to to lack of sequencing coverage. Mendelian (in)consistency can not be verified.

HapIso was able to detect 921 genes with both haplotypes expressed among 9,000 expressed genes. We observed 4,140 heterozygous loci corresponding to position with non-identical alleles among inferred haplotypes. 53% of detected SNVs follow Mendelian inheritance. The number of variants with Mendelian inconsistencies accounts for 10% of the heterozygous SNVs. The remaining SNVs are ambiguous and the Mendelian consistency cannot be verified.

Additionally we check the number of recombinations in the inferred haplotypes. Our approach can theoretically identify recombinations in the transmitted haplotypes. Crossovers between the parental haplotypes result in recombination events in the child’s haplotypes (Figure 4.B). Since recombination events are rare, most of the time they manifest switching errors in phasing. Single-molecule reads are long enough to avoid switching errors, which are confirmed by lack of recombination events, observed in the reconstructed haplotypes.

**Fig. 4.**
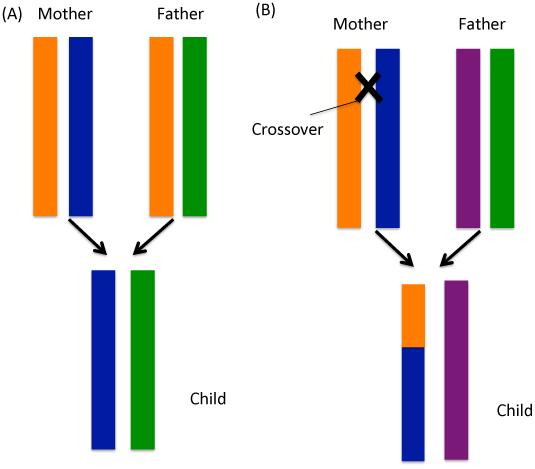
Gene inheritance model. (A) Each parent provides one copy of the gene to a child; child inherits blue haplotype from mother and a green haplotype from father. (B) Haplotypes of mother pair up with each other and exchange the segments of their genetic material to form recombinant haplotype (recombination of orange and blue haplotypes). The child inherits the recombined haplotype. Haplotype from father is inherited with no recombination.

### 3.2 SNV Discovery and Cross Platform Validation

The single-molecule RNA-Seq was complemented by 101-bp paired-end RNA-Seq data of the child. Short RNA-seq reads are used for cross-platform validation of the detected SNVs. The haplotypes assembled by the HapIso were scanned with the 10x coverage threshold to detect the heterozygous loci passed for the validation. The short RNA-Seq reads were mapped onto the hg19 reference genome complemented by the gene annotations (tophat 2.0.13, GRCh37 ENSEMBL). GATK [11] variant caller was used to call the SNVs from the short RNA-seq reads following the publicly available best practices. Additionally, the public catalogue of variant sites (dbSNP), which contains approximately 11 million SNVs, was used to validate genomic position identified as SNVs by single-molecule and short read protocols.

We compared genomic positions identified as SNVs from long single-molecule reads and short reads (Figure 5). First, we compared the positions identified as SNVs by both platforms. 279 genomic positions were reported as SNV by both platforms. Those positions were also confirmed by dbSNR Of those SNVs, 94% are concordant between the platform i.e. contain identical alleles. Among the detected SNVs by the single-molecule protocols, 23 positions are identified as SNVs only by the single-molecule protocol. We investigated the coverage of those SNVs by the short reads. Those SNVs are covered by the short reads with the alternative allele expression under the SNV calling threshold, while the remaining SNVs are not covered by short reads.

**Fig. 5.**
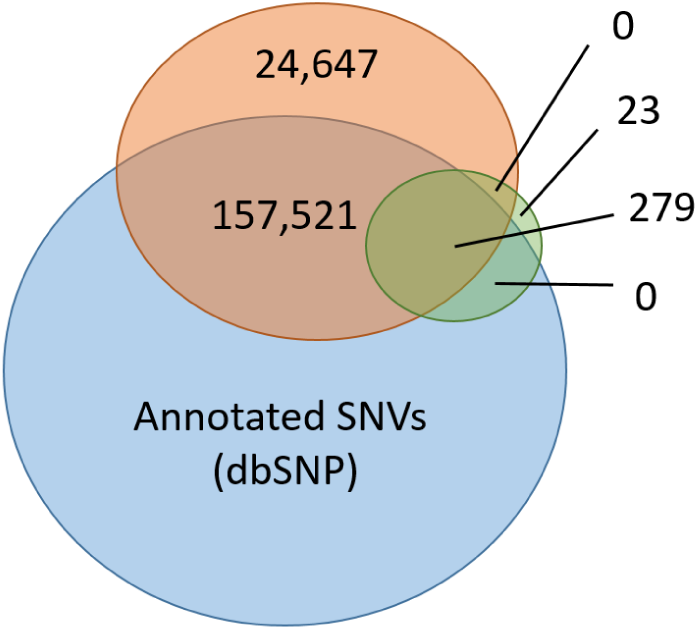
Venn diagram showing the numbers of genomic position identified as SNVs from long single-molecule reads (green) and short reads (orange). SNVs calls from both platforms were match against the dbSNP catalogue of variant sites (blue).

We compared haplotypes assembled from long and short RNA-Seq reads (child sample, GM12878). We use HapCUT [1] to assemble the haplotypes from the short RNA-Seq reads. HapCUT is a max-cut based algorithm for haplotype assembly from the two chromosomes of an individual. GATK is used to generate vcf file with genomic variants required by HapCUT. HapCUT produces multiple contigs per gene shorter than the transcript isoforms, thus limiting the possibility to access haplotype-specific isoforms.

Unfortunately we could not compare our method with HapCUT for the long single-molecule reads. HapCUT is originally designed for the short reads. We are not able to generate the genomic variants (vcf format) required by HapCUT. The GATK tool doesn’t have the best practice pipeline for the Pac Bio RNA-Seq reads.

## 4 Discussion

RNA molecules represent an essential piece of the cell identity, playing an important role as a messenger and regulatory molecule [13]. Long single-molecule protocols provide an unprecedented allele-specific view of the haploid transcriptome. Partitioning the long reads into the parental haplotypes allows us to accurately calculate allele-specific transcript and gene expression and determine imprinted genes [7], [19]. Additionally, it has the potential to improve detection of the effect of cis-and trans-regulatory changes on the gene expression regulation [5], [21]. Long reads allow us to access the genetic variation in the regions previously unreachable by the short read protocols providing new insights into the disease sustainability. Availability of full-length haplotype-specific isoforms opens a wide avenue for the accurate assessment of allelic imbalance to study molecular mechanisms regulating genetic or epigenetic causative variants, and associate expression polymorphisms with the disease susceptibility.

We have presented HapIso, an accurate method for the reconstruction of the haplotype-specific isoforms of a diploid cell. Our method uses the splice mapping and partitions the reads into parental haplotypes. The proposed method is able to tolerate the relatively high error-rate of the single-molecule sequencing and discriminate the reads into the paternal alleles of the isoform transcript. Phasing the reads according to the allele of origin allows efficiently distinguish between the read errors and the true biological mutations. HapIso uses the 2-means clustering algorithm aiming to group the reads into two meaningful clusters maximizing the similarity of the reads within cluster, and minimizing the similarity of the reads from different clusters. Clustering is applied locally for the transcribed regions, which are further reconnected in the segment graph. Each cluster corresponds to the parental haplotype. An error-correction protocol is applied for the reads from the same cluster allowing to correct the errors and reconstruct haplotype-specific isoforms.

Traditional haplotype assembly [1], [12] approaches are designed for whole-genome sequencing and are not well suited for gapped alignment profile offed by the RNA sequencing. Genome-wide haplotype assembly aims to assemble two haplotypes for a chromosome given the collection of sequencing fragments. In contrast, RNA haplotype reconstruction requires to assemble multiple haplotypes of a gene, which each isoform having two parental haplotype copies. ASE detection methods [10] are well suited to determine the allele-specific expression on the on individual variant level further aggregated into gene-level estimates. However, those methods are originally designed for SNV-level read counts and are not applicable to reconstruct full-length haplotype-specific isoforms of a gene.

To our knowledge, our method is the first method able to reconstruct the haplotype-specific isoforms from long single-molecule RNA-seq data. Other approaches [18] quantify the allele-specific expression of the genes using trio family data, while only being able to provide the ratio between allele expression of the genes. Such approaches are not suited to reconstruct the haplotype-specific isoforms and correct sequencing errors. Experimental validation based on the trio family data and orthogonal short read protocol suggests that HapIso is able to tolerate the relatively high error-rate and accurately reconstruct the haplotype-specific isoforms for genes with at least 10x coverage. Deeper sequencing is required to assemble haplotype-specific isoforms of genes with low expression level.

